# Potential Systemic Availability Classification of Chemicals for Safety Assessment

**DOI:** 10.1101/2025.03.04.641374

**Authors:** René Geci, Alicia Paini, Andrew Worth, Lars Kuepfer, Stephan Schaller

## Abstract

The assessment of chemical safety is essential for protecting human health, yet current approaches have severe limitations. They are unable to address the rapidly growing number of substances requiring evaluation and further rely on animal testing, which the public expects to be phased out for ethical reasons. To address this, the European Partnership for Alternative Approaches to Animal Testing (EPAA) proposed the use of a novel framework for the classification of systemic toxicity based on new approach methodologies. One dimension of this framework is the grouping of substances into different classes of Potential Systemic Availability (PSA) concern (low, medium, high). But so far it remained conceptually and practically unclear how this classification can be achieved. Here, we outline the theoretical considerations for a health-protective definition of PSA concern classes and present a method for the quantitative evaluation of this property. Using high-throughput physiologically based kinetic modelling, we are able to classify the PSA concerns of 139 out of 150 EPAA NAM Designathon compounds. Further, we manually annotate these compounds to evaluate the plausibility of predicted classifications against expert judgement. Our results outline under which circumstances it is appropriate to prioritise or deprioritise chemicals due to their toxicokinetic properties. However, we find that most compounds cannot be assessed on their PSA alone and need to be considered medium PSA concern so that classification into any overall systemic toxicity concern class remains possible. Future integration of bioactivity data will be necessary to fully judge the utility of our method and of the entire framework.

## 1. Introduction

The assessment and management of chemical safety is critical for protecting human health. But despite an extensive regulatory framework within the European Union (EU) that has conducted evaluations for decades, the European Environment Agency (EEA) estimates that only about 0.5% of chemicals on the market have been robustly characterised, while approximately 70% lack adequate hazard and exposure data necessary for ensuring their safe use (EEA, 2019). The primary legislation governing industrial chemical risks in the EU is the Registration, Evaluation, Authorisation, and Restriction of Chemicals (REACH) regulation (EU, 2006). But information requirements under REACH are tied to the marketed tonnage of chemicals. For chemicals produced in quantities below 1 tonne per year, no safety information is required. Similarly, for those in the 1-10 tonnes per year category, a basic set of information is required but this is generally insufficient for conducting a comprehensive chemical safety assessment (Botham et al., 2023). Hence, many manufactured chemicals are effectively not subject to any legal act requiring information to ensure their safety. As traditional methods have proven inadequate for addressing the vast number of unassessed chemicals, there is a need for novel approaches to chemical safety assessment.

A major limitation of current safety assessment procedures is their reliance on animal testing. Traditional approaches using animals for chemical evaluation are resource-intensive, making them impractical for assessing the increasing number of substances requiring review. Though historically a standard for toxicological evaluation, animal testing is associated with high costs and lengthy testing timelines. Additionally, there are ethical concerns and societal pressure on policymakers to phase out animal testing. For instance, six out of the ten successful European Union Citizens’ Initiatives have expressed public desire for stronger protection of animals. This further underscores the need for new chemical safety assessment methods that are scientifically robust and safe but also ethically sound.

An essential part of advancing alternative safety assessment procedures is the development of methods that not only predict local but also systemic toxicity, referring to adverse effects that substances have on organs distant from the local site of contact. And while the identification and characterisation of potential hazards are central to toxicity assessment, they need to be complemented by an understanding of a substance’s systemic availability to fully evaluate its potential impact on human health. This requires understanding the toxicokinetics (TK) of substances and how they are absorbed, distributed, metabolised, and excreted (ADME) within the body, as that is an essential component for determining the extent to which substances reach various body parts and whether they pose risks of adverse effects.

The NAM Designathon is an initiative by the European Partnership for Alternative Approaches to Animal Testing (EPAA) that is collaboratively exploring how new approach methodologies (NAMs) may support the assessment of systemic toxicity within a novel regulatory framework not relying on animal testing. At the core of this initiative, a two-dimensional matrix was proposed that integrates toxicodynamic (TD) information about a compound’s bioactivity with its potential TK into an overarching chemical concern classification (Fig. 1A) (Berggren and Worth, 2023). TD and TK properties are considered independently of one another and are also detached from external exposure levels to avoid reliance on exposure data, which is often unavailable or unreliable. It was suggested that under the proposed framework high-concern chemicals could be banned or heavily restricted, medium-concern chemicals could be conditionally used after application-specific risk assessments, and low-concern chemicals could be approved for unrestricted use. The aim of the initiative is to provide levels of protection equivalent to those of traditional safety assessment methods, without attempting to exactly replicate adverse effects observed in animal studies.

**Fig. 1:**
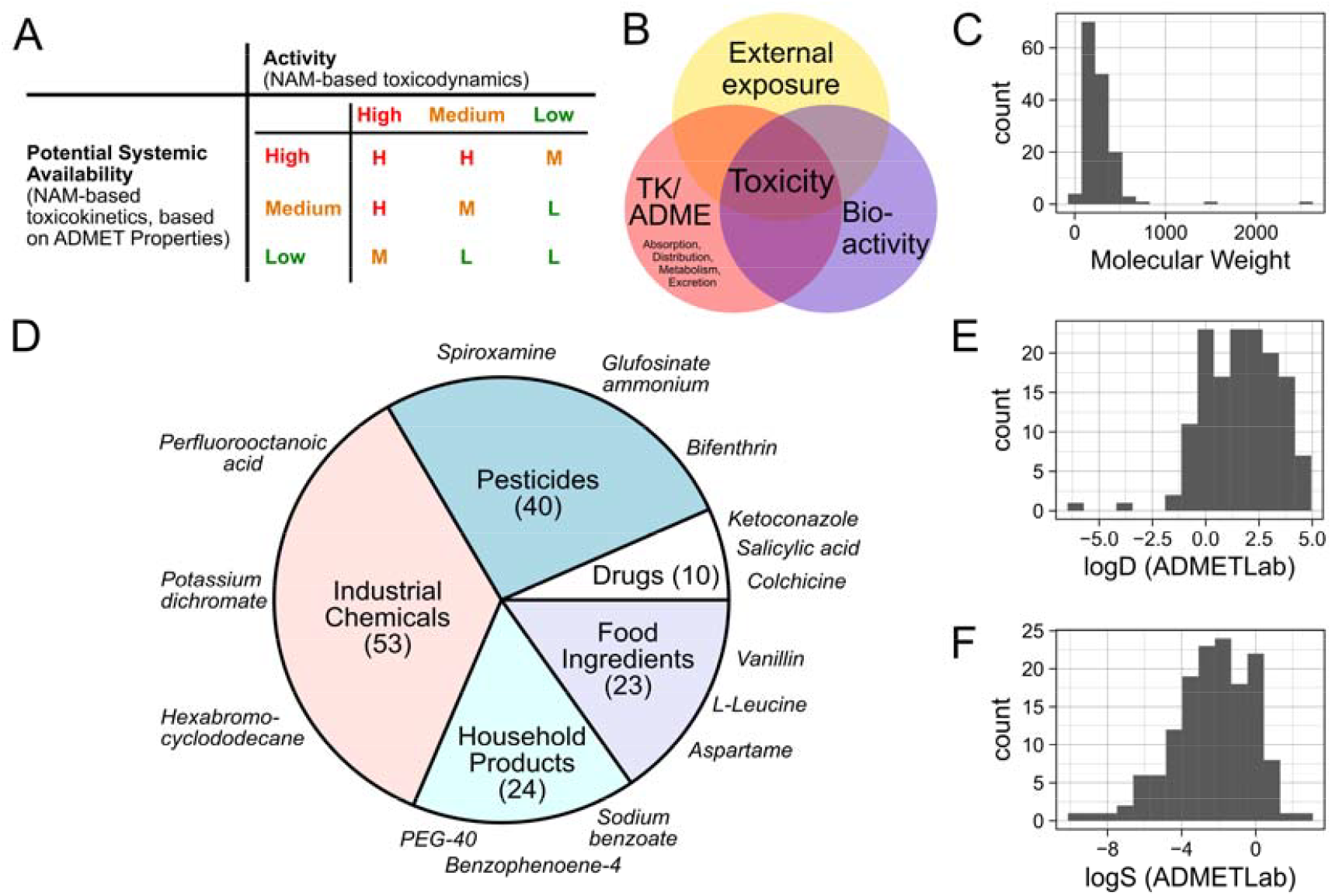
(A) The EPAA NAM Designathon classification matrix integrates Activity and Potential Systemic Availability into an overall systemic toxicity concern class (adapted from Berggren and Worth, 2023). (B) Scheme illustrating that toxicity is determined by external exposure, bioactivity, and toxicokinetics. (C, E, F) Histograms showing the molecular weight, and predicted logD and logS values of NAM Designathon compounds. (D) Distribution of NAM Designathon compounds among different use classes, with three exemplary compounds of each class.

**Fig. 2:**
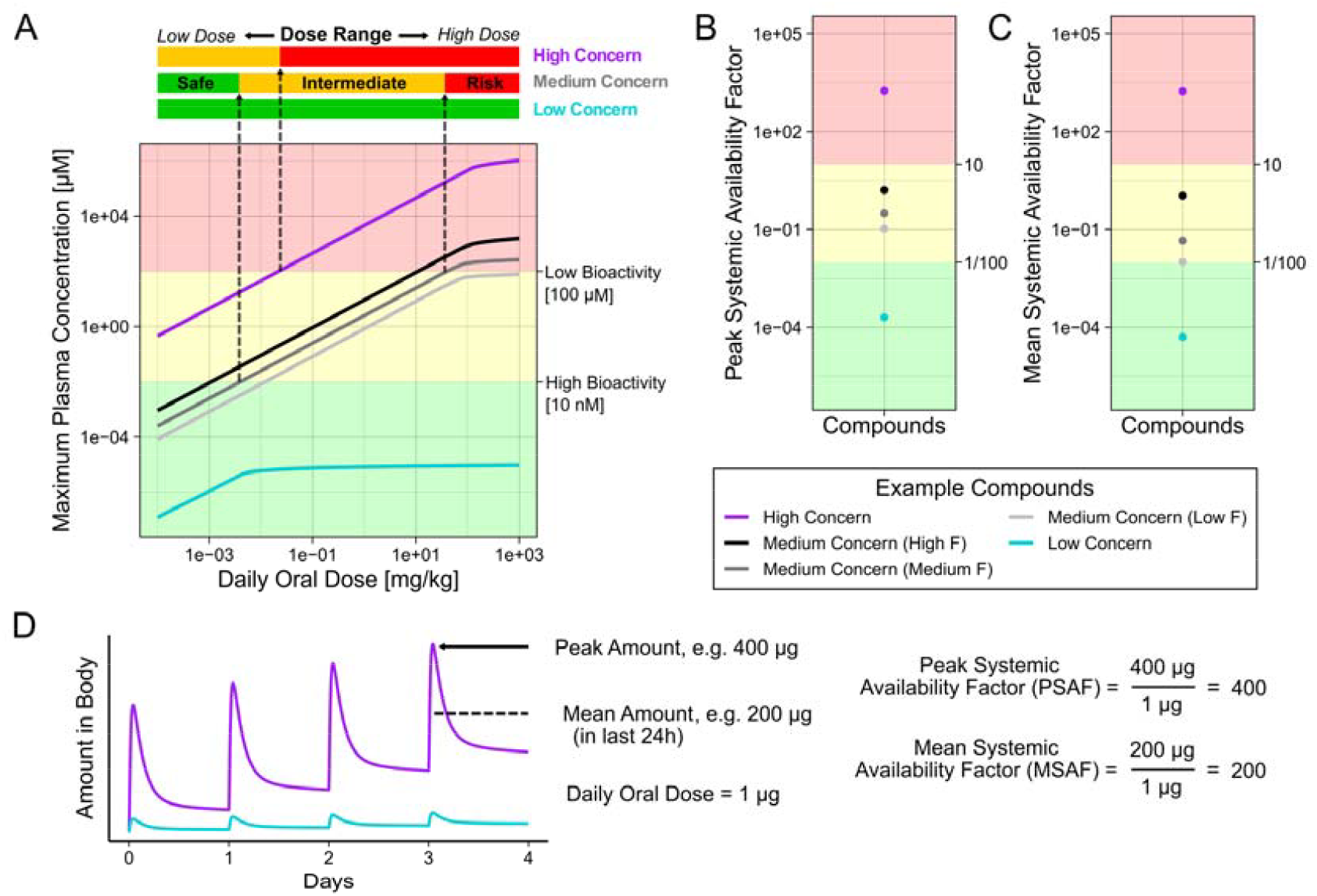
Visual representation for the rationale behind distinguishing different Potential Systemic Availability concern classes. (A) Simulated maximum plasma concentration (Cmax) against oral doses after daily administration for five years of five example compounds as described in Table 1. Marked as safe is the dose range for which the corresponding Cmax value remains below the high bioactivity concentration (10 nM). Risk is the dose range where the Cmax exceeds the low bioactivity threshold (100 µM), and intermediate is the area in between. (B) and (C) show the calculated Peak Systemic Availability and Mean Systemic Availability Factors of the five example compounds, respectively. (D) Example of Peak and Mean Amounts derived from a time-course profile, and calculation of the corresponding PSAF and MSAF values.

To challenge and explore the practical use of this framework, a list of 150 diverse chemicals was proposed by the EPAA, equally representing the three overall systemic toxicity concern classes, and participants were tasked with classifying the substances using NAMs (SI-XXX). The benchmark compounds further represent different use classes, such as industrial chemicals, pesticides, drugs, or substances in household products or in food (Fig. 1C-E). Except for two, all of them are small molecules.

Although the rationale behind the consideration of Potential Systemic Availability (PSA) within the NAM Designathon’s assessment framework was clear, a concrete method for its evaluation had yet to be defined. Toxicity is not only influenced by a chemical’s toxicokinetics but also by external exposure levels and its bioactivity (Fig. 1B). Hence, it is challenging to define a health-protective PSA classification approach that can be used independently, in the absence of information on the other toxicity determining factors. The aim of this work is to explore the scientific rationale for the classification of the PSA of chemicals into three concern classes, as it was suggested for the NAM Designathon. We propose a definition of the different PSA concern classes and a practical method for efficient but also health-protective classification. Additionally, we validate our method by comparing the results against expert judgment-based classifications.

## 2. Methods

### 2.1 Simulation of Maximum Plasma Concentrations of Five Example Compounds

For the simulation of five example compounds, identical physiologically based kinetic (PBK) models using a generic hepatic clearance process and the PK-Sim Standard partitioning and permeability methods were generated in PK-Sim version 11.1.137 (Lippert et al., 2019; Willmann et al., 2003). For simplicity, it was assumed that all five example compounds have identical molecular weights of 300 g/mol, lipophilicity of 1 and fraction unbound values of 50%. Solubility, intestinal permeability and clearance values of compounds differed as presented in Table 1. Then, daily oral administration of solutions of varying doses (1×10^−4^ to 1×10^3^ mg/kg) were simulated for 5 years, and the peak plasma concentration for each dose was extracted. The oral bioavailability of the example compounds was predicted by use of the corresponding standard functionality of PK-Sim, relying on the generated PBK models.

**Table 1:**
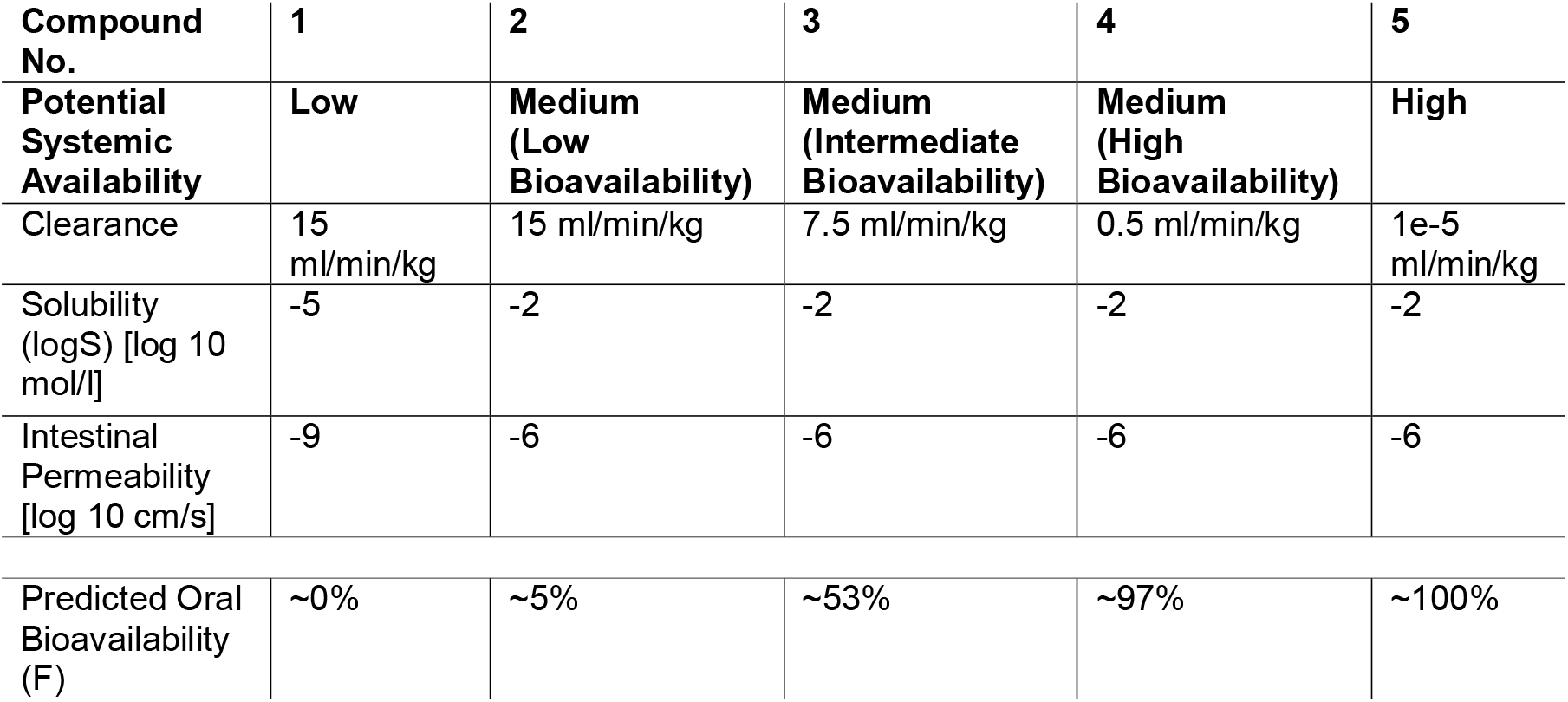
Properties of five example compounds of different Potential Systemic Availability concerns and different oral bioavailability. Other compound properties are assumed to be identical: molecular weight of 300 g/mol, lipophilicity of 1, fraction unbound of 50%.

| Compound No. | 1 | 2 | 3 | 4 | 5 |
| --- | --- | --- | --- | --- | --- |
| Potential Systemic Availability | Low | Medium (Low Bioavailability) | Medium (Intermediate Bioavailability) | Medium (High Bioavailability) | High |
| Clearance | 15 ml/min/kg | 15 ml/min/kg | 7.5 ml/min/kg | 0.5 ml/min/kg | 1e-5 ml/min/kg |
| Solubility (logS) [log 10 mol/l] | -5 | -2 | -2 | -2 | -2 |
| Intestinal Permeability [log 10 cm/s] | -9 | -6 | -6 | -6 | -6 |
| Predicted Oral Bioavailability (F) | ~0% | ~5% | ~53% | ~97% | ~100% |

### 2.2 HT-PBK Modelling of EPAA NAM Designathon Compounds

High-throughput PBK (HT-PBK) simulations of EPAA NAM Designathon compounds were based on previously performed model validations (Gadaleta et al., 2024; Geci et al., 2024). Briefly, all compound properties required for PBK modelling (Kuepfer et al., 2016) were predicted using previously evaluated *in silico* tools: ADMETLab logD, logS and fraction unbound (Fu et al., 2024), ADMET Predictor v12 Fasted State Simulated Intestinal Fluid solubility and intrinsic liver clearance (https://www.simulations-plus.com/software/admetpredictor), OPERA intrinsic liver clearance (Mansouri et al., 2018). For each compound, the default whole-body PBK model implemented in PK-Sim version 11.1.137 was parameterised from R version 4.2.2 (R Core Team, 2022), and administration of 1 µg/kg bodyweight once daily oral solutions were simulated, from which the total amount of compound in the body after five years was extracted as a readout. For partitioning, the methods PK-Sim Standard (Willmann et al., 2005), Rodgers & Rowland (2006) and Berezhkovskiy (2004) were used, and the average of their results was calculated as the final output. Based on these values, Peak Systemic Availability Factor (PSAF) and Mean Systemic Availability Factor (MSAF) values were calculated as shown in equation 1 and 2.

### 2.3 Manual Annotation of EPAA NAM Designathon Compounds

For all 150 EPAA NAM Designathon compounds, we searched for publicly available data that could be informative of their toxicokinetic properties, including *in vitro* data, *in vivo* data from animal and human studies, as well as potentially relevant regulatory classifications such as persistent organic pollutant (POP), persistent bioaccumulative toxic (PBT), very persistent very bioaccumulative (vPvB), and substance of very high concern (SVHC). Further, we extracted basic molecular properties like molecular weight, predicted water solubility and logP values of compounds using above mentioned *in silico* tools.

After having collected potentially relevant data, we decided to classify all endogenous compounds, such as sugars and amino acids, as medium PSA concern. Next, all compounds which possessed TK data and did not show signs of meaningful accumulation were also classified as medium concern. Compounds which did show signs of meaningful accumulation over repeated exposures were classified as high PSA concern. Due to highly heterogenous TK data, e.g., variations in species, administration protocols, investigated tissues, defining absolute criteria for “meaningful” accumulation was not feasible and instead had to determined by experts on a case-by-case-basis holistically considering all available information. Then, in the absence of more information, all remaining compounds without TK data that fell into the “small molecule” space were also classified as medium PSA concern. The two compounds which fell out of this space, both due to large molecular weights above 1500 g/mol, were classified as low concern.

## 3. Results

### 3.1 Definition and Rationale of Potential Systemic Availability Concern Classes

“Potential Systemic Availability” (PSA) is considered an important factor for the assessment of chemical safety in the framework suggested by Berggren and Worth (2023). While the reasoning behind its consideration is clear, the term itself remained underdefined. It was stated that it should encompass toxicokinetics, ADME processes, bioaccumulation, and persistence. However, no official definition exists, nor examples of chemicals representing the different concern categories. To address this gap, we first propose a definition of PSA and justify our definition in direct contrast with a potential alternative.

The definition of PSA concerns we will use in the following is that compounds shall be considered…

> … “*Low PSA Concern*” if their *in vivo* concentrations can be assumed to never reach *in vivo* levels required for bioactivity for almost all relevant exposure scenarios. It can therefore be assumed that there is no exposure dependency for their toxicity, because such compounds would be safe in almost all relevant exposure scenarios.
>
> … “*Medium PSA Concern*” if the reaching of critical *in vivo* concentrations depends on the specific exposure scenario and the bioactivity of the compound. For such compounds, we assume that there will be an exposure dependency for their toxicity, with the compound being safe below a certain exposure threshold but unsafe above it.
>
> … “*High PSA Concern*” if already very small exposures can lead to *in vivo* concentrations required for bioactivity, even if the bioactivity of the compound is only modest. For such compounds, there is no exposure dependency for their toxicity, implying that these compounds would be unsafe in almost all relevant exposure scenarios.

This definition may be understood as a “hard” interpretation of what property the PSA concerns should reflect, as it sets a very high standard for compounds to be classified as either low or high concern. By this definition, most small molecule chemicals, including almost all drugs, pesticides, etc., would fall into the medium PSA concern class, and only extreme chemicals would be considered low or high concern. Further, our definition explicitly refers to peak concentrations *reaching* levels required for *in vivo* bioactivity, which may be transitory. It does not require that these concentrations are also maintained for prolonged periods. In other words, our definition does not rely on exposure over time, as in the area under the curve (AUC), or average concentrations of compounds, but their peak concentration values (Cmax), making this a conservative definition of the term.

Next, we outline a “softer” interpretation of the term to contrast our definition with a potential alternative. This would be to equate PSA with the bioavailability of a compound, which in pharmacology is defined as the fraction of a compound that reaches systemic circulation. In this reading, one may suggest that compounds could be classified as low, medium, or high PSA concern if they have low, medium, or high bioavailability. Whatever the specific thresholds for low, medium and high classification are, it is evident that such an interpretation would lead to a more sensitive division of typical small molecules. Drugs, pesticides, etc., can have drastically varying levels of bioavailability, ranging from just a few percent to full (100%) bioavailability. Hence, when using such a “softer” definition, these compounds would be allocated more evenly into the different PSA concern classes than when using our “hard” interpretation of the term.

To justify why we will continue our work using the “hard” definition, we illustrate *in vivo* plasma concentrations of five example compounds (Table 1). For simplicity and ease of comparability, we assume the five compounds have mostly identical properties, like identical molecular weight, lipophilicity, etc., and only differ in three key parameters: their i) clearance, ii) solubility and iii) intestinal permeability. Using a simple generic PBK model, we then simulate the maximum plasma concentration of these compounds after repeated oral exposure to different once-daily doses for five years. Further, we add two horizontal lines representing thresholds for *in vivo* concentrations of high (10 nM) and low (100 µM) bioactivity for contextualisation. The simulation results (Fig. 1) show that, despite their large differences in bioavailability, compounds 2 – 4 can all reach both the low and high concentration thresholds required for bioactivity. Whether the compounds would do so *in vivo* solely depends on the specific exposure dose and the specific bioactivity of the compounds. Hence, we conclude that these compounds cannot *a priori* be assumed to be safe or unsafe, at least not just based on their PSA, as a “softer” definition would suggest.

However, the situation is different for compounds 1 and 5. Due to its oral absorption being permeability- and solubility-limited, compound 1 would indeed never be able to even reach the 10 nM (“very bioactive”) concentration threshold, regardless of what its daily oral exposure is. Hence, we consider it to be justified to *a priori* classify such a compound as being of generally lower concern. Compound 5 would possess equal *in vivo* concentrations as compound 4 after single oral exposure (SI-Fig. 1), but due to its ability to bioaccumulate over time, it would reach much higher concentrations after prolonged periods of repeated exposure, even if the daily doses themselves are low. In the case of compound 5, this means that already at relatively modest exposures of 0.1 mg/kg, it would reach *in vivo* concentrations above the 100 µM concentration threshold required for bioactivity, even though it is not very bioactive. Due to the increased risk of such compounds to reach concentrations required to cause adverse effects *in vivo*, and the undesirability of the bioaccumulation potential itself, we consider it warranted to *a priori* classify such compounds as being of higher concern.

### 3.2 PBK Model-Based Quantification of Potential Systemic Availability

Having clarified the definition of the PSA concern classes, we have a conceptually more rigorous foundation for the assessment of chemicals. However, from a practical perspective, the question of how this property can be evaluated, ideally efficiently for large numbers of chemicals, remains open.

Three considerations seem particularly important:

i. The method should be organ-agnostic, meaning it should not only rely on concentrations in a single organ. While for illustrative purposes Fig. 1A gives the simplified impression that compounds only have a single *in vivo* concentration, the reality is more complex. The human body is not one homogenous compartment, but it is made up of multiple organs and compound concentrations between those organs can vary substantially. Typically, one would not *a priori* know which organs are potential targets of toxicity and which organ’s concentration would be relevant for safety considerations. Hence, selecting a single organ for classification purposes is not trivial. It may even be counterproductive. For example, many in the body accumulating chemicals are highly lipophilic and, therefore, exhibit higher concentrations in fatty tissues, and, conversely, lower concentrations in plasma. If one only tracks concentrations in plasma, for example, then a more lipophilic, and for this reason bioaccumulating compound, may falsely appear less concerning than a less lipophilic compound.
ii. To correctly capture the drastic differences in compounds’ potential to bioaccumulate, it is required to consider repeated exposures over longer time periods, not just short-term or even single exposures.
iii. The output should reflect a general, intrinsic compound property that is ideally independent of arbitrary assumptions or threshold values, e.g. specific exposure doses or concentration thresholds. Such arbitrary, or at least subjective, values can be controversial and may undermine the general acceptance of assessments.

Based on the outlined considerations, we define the Peak Systemic Availability Factor (PSAF) as

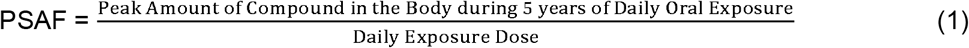

This metric can be understood as the peak total body burden of a compound relative to its daily exposure, and it satisfies all three outlined requirements: (i) It considers the total amount of compound in the body, not just in arbitrarily selected organs. (ii) It considers accumulation after repeated exposure. (iii) The value itself is unitless and reflects an intrinsic compound property. Fig. 1B shows that the PSAF quantitatively captures the expected differences between the five example compounds: Compound 1 has the lowest PSAF of 2×10^−4^, compounds 2 – 4 fall between 0.1 to 1.6, and compound 5 has a PSAF of 1789. Fig. 1D gives a visual representation of how this parameter would be derived from a specific time-course profile.

Despite avoiding arbitrary assumptions for the calculation of the PSAF itself, its translation from numerical values into the three PSA concern classes does eventually necessitate the definition of thresholds, which, as outlined before, is arbitrary. We here decide on a value of 10 for the distinction between medium/high concern and 1/100 for the distinction between low/medium. Practically, this means that a compound is considered of high concern if it can reach 10 times higher amounts in the body than the daily oral exposure to that compound is. Conversely, it would be of low concern if the total amount of compound in the body is never higher than 1% of the daily oral exposure.

To explore and contrast the PSAF for scientific purposes with a second potential evaluation metric, we further define the Mean Systemic Availability Factor (MSAF) as

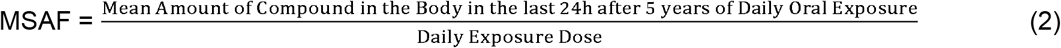

The MSAF is very similar to the PSAF, only that it does not evaluate the highest (peak) amounts of compound in the body, but its time-averaged mean amount. Similar to bioavailability, this makes the MSAF more sensitive to distinguishing chemicals than the PSAF. Compounds that are being cleared more rapidly can potentially have similar PSAF values as compounds that are cleared only slowly, but their MSAF values would be lower. For ease of comparison, we keep the same classification thresholds of 10 and 1/100, although, there may be good reasons to use different thresholds for the two metrics. Fig. 1C shows that the five example compounds can also be classified correctly by use of the MSAF. Although, the MSAF values of the three medium-concern compounds lay closer to the low-concern compound than it is the case for the PSAF values.

### 3.3 Plausibility Evaluation of Potential Systemic Availability Classifications

To demonstrate the utility of our classification metrics, we calculate PSAF and MSAF values for a large set of diverse substances, the 150 EPAA NAM Designathon compounds. For this, we use a previously described high-throughput PBK modelling approach (Geci et al., 2024), which enables the rapid generation of generic compound PBK models. This allows the high-throughput prediction of *in vivo* concentrations, and hence the efficient evaluation of our proposed scores for a large dataset. Nonetheless, we would like to highlight that our method can be used with any type of PBK model and is independent of how the used PBK models are parameterised.

HT-PBK modelling-based predictions could be made for 139 out of the 150 EPAA NAM Designathon compounds. The remaining 11 compounds were inorganics, metals or organometallics, for which the input *in silico* tools could not provide the required property predictions. Predicted PSAF and MSAF values of the EPAA NAM Designathon compounds are shown in Fig. 3A and 4A, respectively. Only 2 out of the 139 successfully predicted compounds have a PSAF < 1%, and hence are classified as low PSA concern. 16 compounds were predicted to have a PSAF > 10 and end up in the high concern category. The remaining 121 compounds all have PSAF values between 1% and 10 and therefore are classified medium PSA concern. Using MSAF generally leads to similar classification results as PSAF, only that more compounds are considered low PSA concern (20 instead of 2). The number of high PSA concern compounds is similar to PSAF (14 instead of 16). As expected, we observe that MSAF values are distributed more widely than PSAF values.

**Fig. 3:**
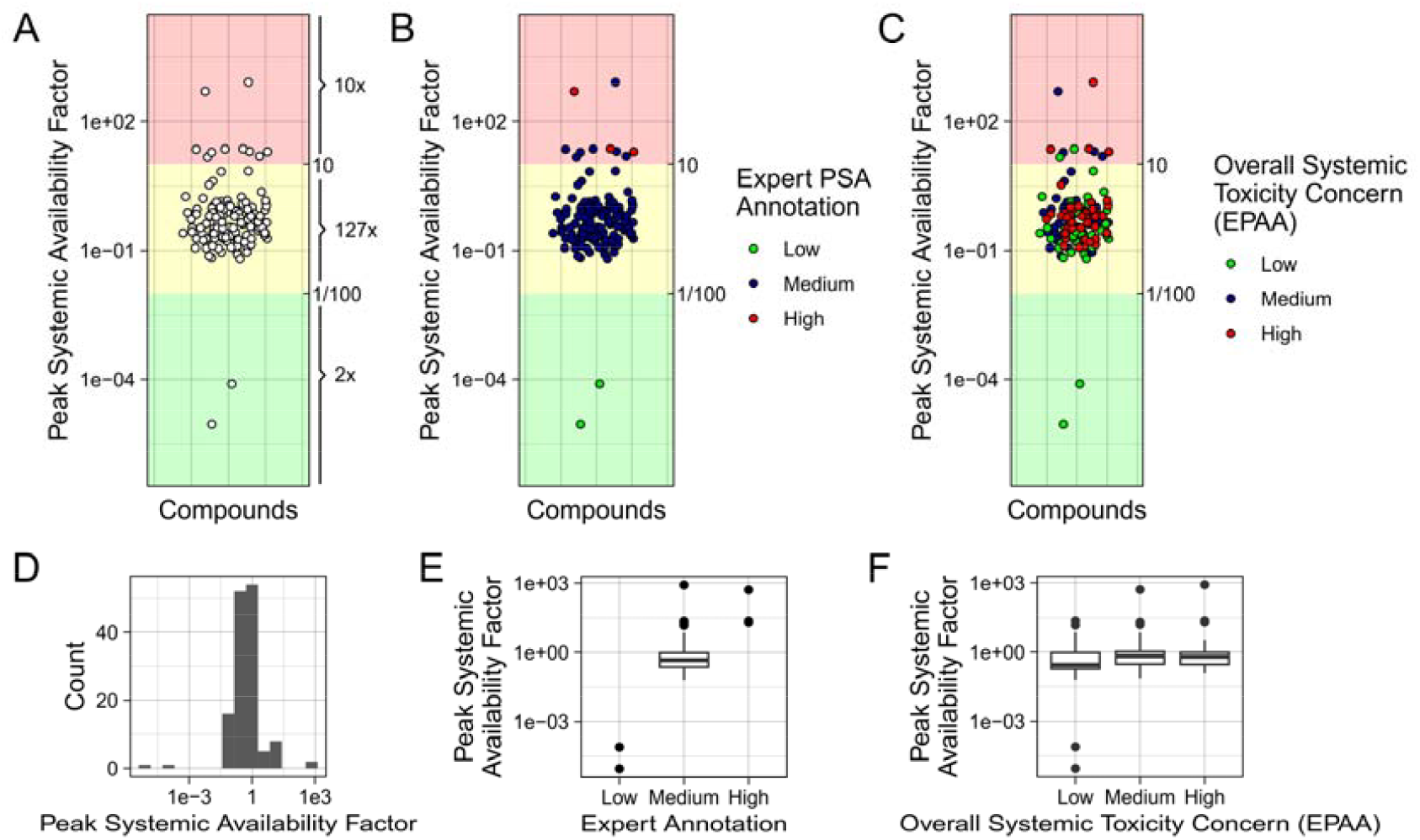
Peak Systemic Availability Factors of the 139 EPAA Designathon Compounds for which HT-PBK predictions could be made. (A,D) Distribution of PSAF values with jittered x-axis. (B) Coloured by manually annotated reference classification. (C) Coloured by overall systemic toxicity concern given by EPAA. (E, F) Comparison of PSAF values of the different manually annotated Potential Systemic Availability or overall concern classes.

**Fig. 4:**
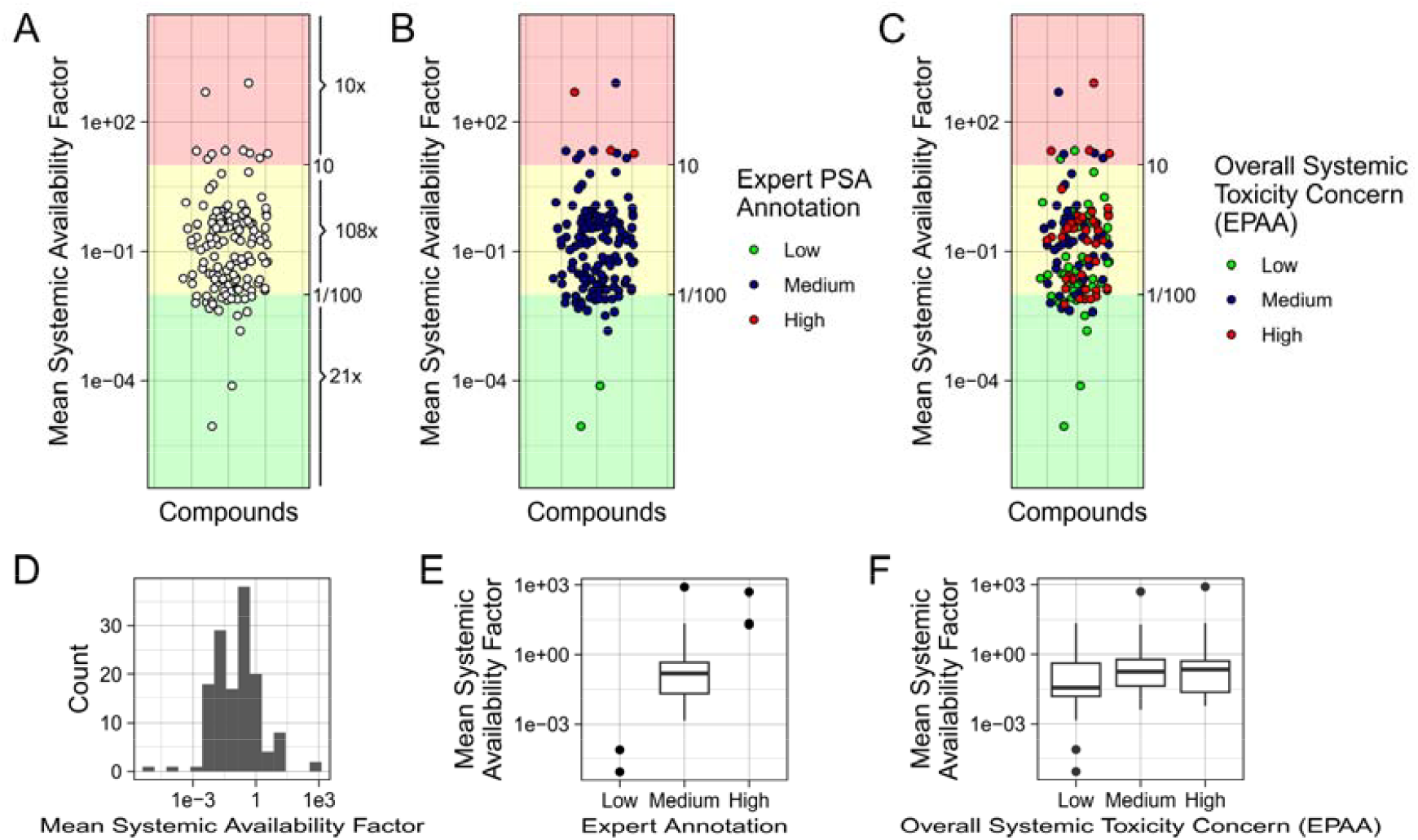
Mean Systemic Availability Factors of the 139 EPAA Designathon Compounds for which HT-PBK predictions could be made. (A,D) Distribution of MSAF values with jittered x-axis. (B) Coloured by manually annotated reference classification. (C) Coloured by overall systemic toxicity compound concern given by EPAA. (E, F) Comparison of MSAF values of the different manually annotated Potential Systemic Availability or overall concern classes.

To have a comparison for assessing the plausibility of our predicted classifications, we manually established a reference classification based on review of public data and expert judgment of the NAM Designathon compounds. The details of this process are outlined in Methods 3.3, and the manually annotated reference classifications are provided in SI-XXX. It proved challenging to annotate compounds since the type, quality and consistency of data available for them were highly variable. For some compounds, high-quality *in vivo* data after repeated oral exposure in humans was available, while others only had lower-quality data from animal or *in vitro* studies, or not even any TK-relevant information at all.

We classified all endogenous compounds (amino acids, sugars, fatty acids) as medium PSA concern, assuming such endogenous substances would be absorbed rather efficiently, yet also not bioaccumulate. One potential exception to this was Ergocalciferol (Vitamin D2), for which a classification as high PSA concern was considered because it is a fat-soluble vitamin which can accumulate in the body over time. But since its half-life is only on the order of days, we eventually decided that its accumulation potential was still not high enough and hence also classified it as only medium PSA concern.

All other compounds with TK data available had been reported to be absorbed orally to some extent, at least so much that for none of them classification as low PSA concern seemed justified. Their reported clearances, half-lives or potential to accumulate in tissues, however, varied quite substantially. Compounds that had been reported to bioaccumulate strongly or have half-lives of at least weeks were then investigated more closely for potential classification as high PSA concern. We eventually classified three compounds as high PSA concern (Perfluorooctanoic acid, Perfluoroheptanoic acid, Hexabromocyclododecane), and the remaining ones as medium. We observed that two compounds (Polyoxyl(40)stearate and

2-Hydroxypropyl-beta-cyclodextrin) possessed very large molecular weight (> 1500 g/mol) and classified both as low concern, assuming they are not absorbed orally to any relevant extent. All remaining compounds for which we could not find specific TK data fell into the small molecule domain with less than 1000 g/mol molecular weight. In the absence of further evidence, we hence assumed these are all medium PSA concern. For some compounds, there was additional uncertainty in their correct reference classification due to special properties like depuration by evaporation (Decamethylcyclopentasiloxane; (Gobas et al., 2015) or potential explosiveness (Dipicrylamine). Eventually, we classified 2 of the compounds as low, 134 as medium, and 3 as high PSA concern.

Comparing our HT-PBK predictions of PSAF values to the expert judgement-based reference classifications reveals that the bulk of compounds had been classified correctly, with 132 out of 139 chemicals grouped identically (Table 2). 7 compounds that were manually classified as only medium PSA concern, were predicted to be of high concern by HT-PBK predictions. However, this type of false-positive misprediction is rather unproblematic for assessment purposes. No compound was predicted to have lower classification than decided by expert judgment. For MSAF-based classifications, however, we do find some compounds being falsely predicted to have low classification, while experts classified them medium PSA concern (SI-Table 2). Even after investigating these compounds in more depth, we did not find any evidence suggesting that these might be of low concern.

**Table 2:**
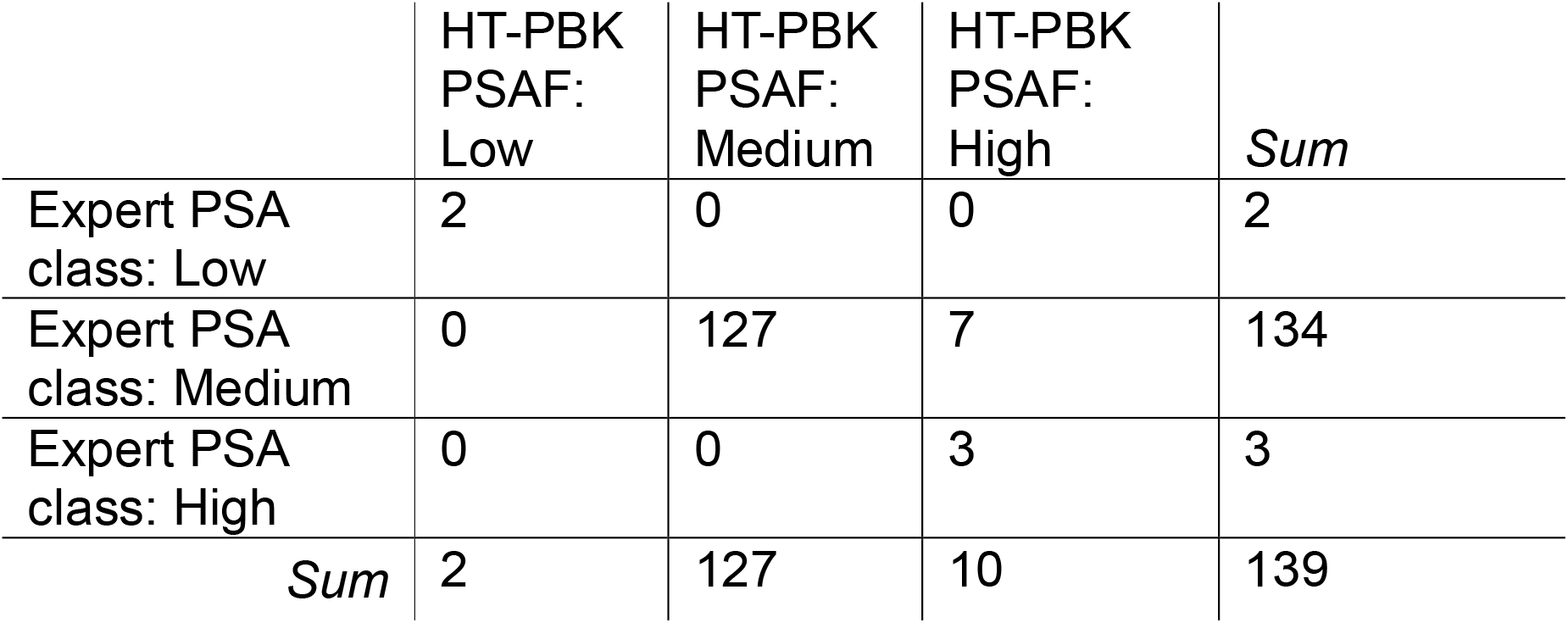
Comparison of classifications of 139 EPAA Designathon Compounds based on HT-PBK predictions using the Peak Systemic Availability Factor (PSAF) against expert judged classification of Potential Systemic Availability (PSA).

|  | HT-PBK<br>PSAF:<br>Low | HT-PBK<br>PSAF:<br>Medium | HT-PBK<br>PSAF:<br>High | *Sum* |
| --- | --- | --- | --- | --- |
| Expert PSA<br>class: Low | 2 | 0 | 0 | 2 |
| Expert PSA<br>class: Medium | 0 | 127 | 7 | 134 |
| Expert PSA<br>class: High | 0 | 0 | 3 | 3 |
| *Sum* | 2 | 127 | 10 | 139 |

Eventually, we also compared predicted PSA classifications to the overall systemic toxicity concern provided by the EPAA. The overall concern categorisation considers not only the PSA but also the bioactivity of a substance and was derived using traditional toxicity data. Hence, these classifications are not expected to be identical to our PSA classifications. But since PSA is one of the two relevant factors of the overall systemic toxicity concern, the comparison may still give a rough indication of the plausibility of our PSA classifications. At least, there should be no compound of overall high concern that is only classified low PSA, and no compound that is of overall low concern but classified high PSA. Indeed, we observe that the first is not the case. The 2 compounds classified as low PSA are also classified low overall concern by the EPAA. However, out of the 16 compounds predicted to be high PSA by use of HT-PBK modelling and the PSAF, 3 compounds are classified only low overall concern. Assuming the overall concern classification by the EPAA is correct, this suggests that the HT-PBK based classification of these compounds is too high, and that these are false positives. Notable, we observe that there is a general tendency of low overall concern compounds to have lower PSAF and MSAF values. This effect is more pronounced for MSAF values, although there is still large overlap between the three overall concern groups. We also observe that, unlike for the PSAF, some high overall concern compounds have only low MSAF values. Hence, it seems that the more strongly spread distribution of MSAF values, and its potential ability to better distinguish lower concern compounds comes at the risk of causing more false-negative classifications. Only the more conservative peak-based classification led to no false-negative classifications.

## 4. Discussion

We here proposed a definition of PSA concern classes, and a method for quantitatively measuring this property. Based on expert judgement, we manually annotated a benchmark dataset of diverse chemicals from the EPAA NAM Designathon and compared our predicted classifications to this annotation. This demonstrated that HT-PBK modelling could be used to correctly predict chemical PSA classifications in a high-throughput manner.

The conceptual rationale for our classification of compounds directly builds on the fundamental dogma of toxicology that chemicals can only cause adverse effects if their doses, and hence their *in vivo* concentrations, are sufficiently high (“the dose makes the poison”). We show that certain compounds can be assumed to be unlikely to reach such *in vivo* concentrations, and we argue that therefore they deserve to be treated more permissively in chemical safety assessment. Conversely, we showed that compounds that have the potential to accumulate in the body over time can reach harmful *in vivo* concentrations already at low daily doses, even if they only possess modest bioactivity. Hence, those deserve to be treated more cautiously.

Further, the potential for bioaccumulation comes with additional implications that are undesirable from a safety perspective. For instance, if the bioactivity of a bioaccumulating substance is underestimated, there may be a significant delay between the exposure to chemicals and the observation of their toxic effects, as it may take years for substances to accumulate to steady-state levels in humans. Subsequently, their adverse effects may only become apparent long after widespread exposure. By this time, these chemicals may have already been used and widely distributed, making retroactive mitigation and response efforts highly challenging. This is currently a major problem regarding per- and polyfluoroalkyl substances (PFAS; (Brunn et al., 2023; Sunderland et al., 2019).

Our definition of PSA reflects a health-protective interpretation of the term as it assumes that reaching peak levels once may be sufficient to trigger adverse effects *in vivo*. However, evidence suggests that for some chemicals, toxicity is more closely linked to exposure over time (AUC), rather than peak (Cmax) levels (Groothuis et al., 2015; Tzimas et al., 1997). This raises the possibility that AUC may be a better metric for the classification of chemicals. To explore this possibility, we investigated whether a time-average-based approach is useful for classification purposes. Our findings demonstrate that the most straightforward approach using bioavailability as a proxy for PSA, is not suitable for classification. Instead, we proposed a method that accounts for exposure across all organs and considers accumulation from repeated exposures over time.

For our benchmark chemical set, we find that peak (PSAF) and time-averaged (MSAF) classifications yield comparable results, with most NAM Designathon compounds clustering around moderate values with only a few extremes. Hence, both metrics could potentially be used for the classification of chemicals. Further, we observe that compounds of low overall systemic toxicity concern tend to also have lower MSAF values, and the distinction between those and higher overall concern compounds is more pronounced with MSAF than with the PSAF metric. This seems reasonable, as compounds that exhibit lower exposure have, all else being equal, less potential for causing adverse effects.

However, we also find exceptions to this trend, where high overall concern compounds display low MSAF values. We speculate that those represent cases where chemical toxicity is driven by peak levels, rather than exposure over time. For instance, phosphamidon is an organophosphate pesticide, which are cleared relatively efficiently (Moerenhout et al., 2024), and MSAF-based classification also suggests it is of only low PSA concern. In the context of the EPAA Designathon, it is classified as overall high systemic toxicity concern. And indeed, phosphamidon is a known cholinesterase inhibitor that causes adverse effects above certain concentration thresholds (Sidhu et al., 2019). Therefore, it seems plausible that this compound’s toxicity is driven by Cmax, rather than AUC, and that it is only classified appropriately by consideration of peak levels, but not by time-averaged levels. Given that it is typically unknown beforehand whether peak concentration or exposure over time is the primary driver of toxicity for a given compound, we conclude that a peak-based approach is better suited for safety assessments, as it is the more conservative metric and avoids false-negative classifications, which may arise when using time-average-based values. Even though false-negative classifications may be rare, they can undermine public and regulatory acceptance.

Further, we manually annotated the PSA classes of the Designathon compounds to have a reference for the evaluation of our predicted classifications. Consistent with our predictions, we concluded that most compounds are best classified as medium PSA concern. These compounds, therefore, cannot *a priori* be considered more or less safe, since they may end up in any of the three systemic toxicity concern categories, depending on their specific bioactivity. Considering that most of the investigated compounds are small molecules, like endogenous metabolites, drugs, and pesticides, this seems plausible. These substances have been deliberately selected by nature or industry to be bioavailable and/or exert bioactive effects. On the other hand, this observation is perhaps counterintuitive, as the NAM Designathon compounds were selected to equally represent the three overall systemic toxicity concern classes, with 50 compounds from each class. Hence, one might have expected them to be distributed evenly across the three PSA concern classes.

It is important to note that the data available for establishing reference classifications was highly heterogeneous, making it challenging to confidently assign reference classes to many compounds. We acknowledge the subjectivity of our manual annotation process, which relied on interpreting heterogeneous data on a case-by-case-basis, without established standards or protocols. Nevertheless, we believe the general trends described here are robust. Alternative data sources, such as human biomonitoring data, may provide insights for more accurate classification of some chemicals, although such an analysis was beyond the scope of this study.

Because we evaluated PSAF and MSAF values by using HT-PBK modelling, our results are subject to the general limitations of this method. Primarily, the accuracy of our predictions depend on the reliability and applicability of the underlying *in silico* tools providing the input parameters for PBK models. When input parameters are incorrect, the simulation outputs will also be flawed. Addressing these limitations requires a sustained effort to further validate *in silico* tools, to expand their applicability domains, and to improve their reliability and accuracy. For the shown application, it is especially relevant that *in silico* tools not just accurately predict average values but also extreme outliers, such as compounds with exceptionally low or high clearance, fraction unbound, or intestinal permeability values.

These extreme properties are often the primary drivers for classification into low or high PSA concern classes. Unfortunately, *in silico* tools currently struggle precisely with this, since their training datasets often underrepresent compounds having extreme values (Cherkasov et al., 2014; Kirchmair et al., 2015). Addressing this will be essential for improving the accuracy of *in silico* classifications like those presented here. Alternatively, our PBK models may be parameterised using *in vitro* data instead. However, *in vitro* methods also possess biases and may potentially face similar difficulties in handling extreme chemicals (Houston and Galetin, 2003; Proença et al., 2021).

Limitations of the *in silico* tools used here also explain why certain groups of substances, like metals or inorganics, could not be assessed with our approach. Nevertheless, the coverage of 139 out of 150 compounds of interest was high, suggesting this may not be typically problematic for applications in safety assessment. Additionally, we identified some non-standard processes that may influence compounds’ PSA concern but were not included in our generic PBK models. Examples include the depuration of volatile compounds by breathing, chemical instability, gut metabolism, and biliary clearance. Ideally, all relevant processes should be included in PBK models to enable comprehensive and reliable evaluations. However, such detailed modelling was beyond the scope of our analysis and the development of predictive tools for these processes remains an ongoing challenge. The same applies to the prediction of transporter activities, which can significantly affect local organ concentrations and sometimes may be a crucial factor for *in vivo* toxicity (Klaassen and Aleksunes, 2010; Najjar et al., 2022). Finally, a comprehensive assessment of exposure should encompass multiple exposure routes, including, for instance, dermal and inhalation exposure. Here, we focused solely on oral absorption, as it is one highly relevant route of exposure. Nevertheless, our approach can also be applied to other exposure routes.

An alternative to the PBK modelling-based approach is to use rule-based classification schemes or simple regression models relying on selected chemical properties. This is, for example, done for predicting bioaccumulation in aquatic organisms and bioconcentration factors (Arnot and Gobas, 2006). However, this requires the subjective definition of classification thresholds and the selection of individual compound properties to be considered (Arnot et al., 2010). In contrast, PBK modelling possess the power to integrate various relevant molecular properties that jointly determine quantitative *in vivo* behaviours. It is therefore better suited to capture quantitative *in vivo* outcomes and the complex relationships between the various molecular properties. This is why PBK modelling is increasingly applied to evaluate *in vivo* exposure to chemicals (Coecke et al., 2013; Tonnelier et al., 2012; Wambaugh et al., 2015; Wetmore et al., 2012), although ultimately the choice of method may depend on the context, available information and the required accuracy.

We observed for both predicted and manually annotated classifications that most compounds should be considered medium PSA concern, yet their overall systemic toxicity concerns are distributed equally among the three classes. This suggests that the overall concern of compounds is driven mainly by differences in compound bioactivity, which questions the utility of TK-based classifications. Regarding this, we would like to point out two things.

Firstly, identifying high PSA concern compounds is valuable, even if they only represent a small subgroup of chemicals, because it enables prioritised assessments of such compounds. This is a signification benefit given the existence of thousands of untested chemicals on the market. Secondly, when considering the broader chemical universe, it is conceivable that a substantial number of chemicals may fall into the low PSA category. But, potentially, these chemicals were just not represented in the NAM Designathon dataset, and are not even routinely tested, because i) their extreme chemical properties, such as low permeability, make them difficult to test *in vitro*, and ii) because based on their low bioavailability they are not expected to cause relevant adverse effects that would meet existing classification criteria. Therefore, our classification approach may simply reflect the current handling of such compounds and supports the rationale for their current regulatory treatment. For example, under our definition of PSA, most polymers would likely be classified as low concern, as they are generally regarded to not readily cross biological membranes and hence to exhibit minimal systemic toxicity. This explains why polymers are subject to regulatory exceptions (U.S. Environmental Protection Agency, 1997), aligning well with our results, yet such compounds were not represented in our evaluation dataset.

Finally, the application of the presented method remains to be explored more concretely within the context of a novel framework for regulatory decision-making. It is an ongoing effort to outline how the current system of chemical safety assessment may be changed to finally phase out animal testing (Ball et al., 2022; Berggren and Worth, 2023; Schmeisser et al., 2023; Worth et al., 2024; Worth and Berggren, 2024). We showed that the consideration of TK properties in the form of the PSAF could support an *a priori* prioritisation of certain chemicals, and the de-prioritisation of others. This may be useful for classifying, or at least triaging, compounds within a novel tiered safety assessment framework. Since the here used high-throughput PBK model approach only requires *in silico* predictions of a few compound properties, it allows the rapid classification of the large number of currently unassessed substances. Even if one would use more reliable *in vitro* data for parameterisation, it would still be possible to efficiently conduct such an analysis for large numbers of compounds (Rotroff et al., 2010; Wetmore et al., 2012).

However, we also found that many compounds cannot be (de)prioritised outright, as most were classified into the medium PSA concern category. From this category, classification into any overall systemic toxicity concern class remains possible, depending on the bioactivity of compounds. In cases where the bioactivity of compounds is also neither particularly low nor high, it would be necessary to conduct a more in-depth comparison of their *in vivo* concentrations after specific exposures against their bioactivities to determine bioactivity exposure ratios (BERs) or concrete thresholds for (un)safe exposure levels (Wetmore et al., 2015). Such more elaborate, in-depth evaluations could build directly on the same HT-PBK models used for an initial PSA classification.

In the absence of information on specific bioactivity and internal exposure levels, the presented approach had to be based on highly conservative assumptions. We anticipate that integrating bioactivity data into the analysis will eventually enable a larger number of NAM Designathon compounds to be classified more definitively as either low or high systemic toxicity concern. This will finally enable a comparison of overall systemic toxicity concern classifications using NAMs against classifications based on traditional toxicological studies.

## Abbreviations

ADME: Absorption, Distribution, Metabolism, and Excretion
AUC: Area under the curve
BER: Bioactivity exposure ratio
CLP: Classification, Labelling and Packaging
Cmax: Maximum concentration
EEA: European Environment Agency
EPAA: European Partnership for Alternative Approaches to Animal Testing
EU: European Union
HT-PBK: High-throughput PBK
MSAF: Mean Systemic Availability Factor
NAM: New approach methodology
PBK: Physiologically based kinetic
PBT: Persistent, bioaccumulative, toxic
PFAS: Per- and polyfluoroalkyl substances
POP: Persistent organic pollutant
PSA: Potential Systemic Availability
PSAF: Peak Systemic Availability Factor
REACH: Registration, Evaluation, Authorisation and Restriction of Chemicals
SVHC: Substance of very high concern
TD: Toxicodynamic
TK: Toxicokinetic
vPvB: very persistent, very bioaccumulative

## Data availability

All data is available in the Supplementary Information.

## CRediT Author statement

René Geci: Conceptualization, Data curation, Formal Analysis, Investigation, Methodology, Visualization, Writing – original draft; Alicia Paini: Conceptualization, Data curation, Investigation, Methodology, Supervision, Writing – review & editing; Andrew Worth: Data curation, Investigation, Methodology, Writing – review & editing; Lars Kuepfer: Supervision, Visualization, Writing – review & editing; Stephan Schaller: Funding acquisition, Supervision, Writing – review & editing

## Supplementary Information

**SI-Fig. 1:**
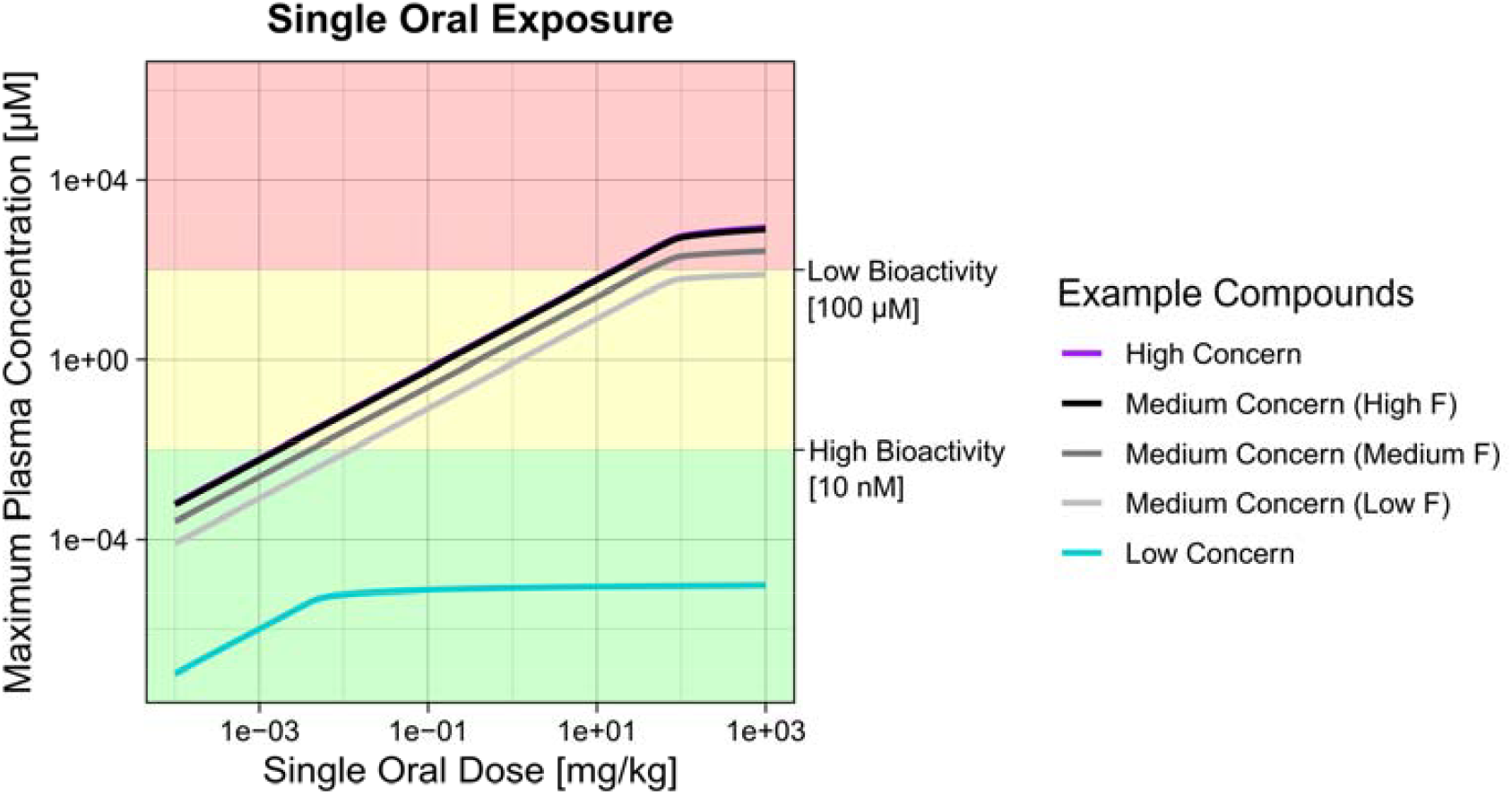
(A) Maximum plasma concentration (Cmax) against oral doses after single administration of five example compounds with varying properties as described in Table 1.

**SI-Table 1:** Comparison of classifications of 139 Designathon Compounds based on HT-PBK predictions using the Mean Systemic Availability Factor (MSAF) expert judged classification of Potential Systemic Availability (PSA).

|  | HT-PBK<br>MSAF:<br>Low | HT-PBK<br>MSAF:<br>Medium | HT-PBK<br>MSAF:<br>High | Sum |
| --- | --- | --- | --- | --- |
| Expert PSA<br>class: Low | 2 | 0 | 0 | 2 |
| Expert PSA<br>class: Medium | 19 | 108 | 7 | 134 |
| Expert PSA<br>class: High | 0 | 0 | 3 | 3 |
| Sum | 21 | 108 | 10 | 139 |

